# Engineering murine GITRL for antibody-mediated delivery to tumor-associated blood vessels

**DOI:** 10.1101/2020.08.26.268375

**Authors:** Jacqueline Mock, Itzel Astiazaran Rascon, Marco Stringhini, Marco Catalano, Dario Neri

## Abstract

Preclinical evidence has suggested that the Glucocorticoid-Induced TNFR-related protein (GITR) may be valuable a target for the development of anticancer therapeutics, but clinical studies with GITR ligand (GITRL) have been disappointing. Here, we report the development of a fusion protein featuring GITR ligand (GITRL) fused to the F8 antibody which targets the alternatively-spliced EDA domain of fibronectin, a tumor-associated antigen often found around the tumor neovasculature. Five different formats for F8-GITRL fusion proteins were cloned and characterized, but quantitative biodistribution studies failed to evidence a preferential accumulation at the tumor site. The *in vivo* tumor targeting properties of F8-GITRL could be substantially improved by enzymatic deglycosylation or site-directed mutagenesis of the *N*-glycosylation consensus sequence. However, therapy studies in a murine model of cancer with the glycoengineered F8-GITRL N74S and N157T variant failed to elicit a durable anti-tumor response, both in monotherapy and in combination with PD-1 blockade.

**HIGHLIGHTS:**

- Different formats of fusion proteins featuring Glucocorticoid-induced TNFR-related protein ligand (GITRL) fused to a tumor-targeting antibody were produced.
- The tumor uptake of the fusion proteins could be increased by enzymatic deglycosylation of the fusion protein or by site-directed mutagenesis of the *N*-glycosylation consensus sequences.
- The fusion protein developed in this study failed to show any anti-tumor activity either alone or in combination with PD-1 inhibition.

## 1. INTRODUCTION

The recent clinical success of immune checkpoint inhibitors in a subset of patients^1–3^ has fueled interest in the development of additional immunostimulatory products for the treatment of cancer^4,5^. Attractive targets include T cell costimulatory receptors such as Glucocorticoid-induced TNFR-related protein (GITR)^6^. GITR is of particular interest since there is evidence that it not only delivers additional costimulatory signals to activated effector T cells and enhances the survival of this subset^7^, but that it also reduces the suppressive function of regulatory T cells^8^. In addition, it was shown to render effector T cells more resistant to suppression by regulatory T cells^9^. It was recently shown that the delivery of agonistic signals to GITR combined with PD-1 inhibition would revert CD8+ T cell dysfunction and enhance the memory function of this subset^10^. While most GITR agonists in preclinical and clinical development are antibodies, the delivery of recombinant soluble GITR ligand (GITRL) is also an option^6^.

One advantage of recombinant soluble ligands is that they are amenable to multimerization by linking several subunits via short oligopeptide linkers and by genetically fusing these single-chain multimers to antibodies or antibody fragments^11,12^. Importantly, clustering of receptors of the TNF superfamily is a prerequisite for signaling through this class of receptors^13,14^. Researchers at Apogenix have developed a fusion protein of a single-chain trimeric GITRL and an Fc portion yielding a hexavalent GITR agonist which showed superior *in vitro* agonistic activity compared to a clinical-grade agonistic antibody and showed some anti-tumor activity in preclinical models of cancer^12^. In spite of these promising preclinical results, a recent clinical trial featuring the use of an Fc fusion of human GITRL failed to show objective responses, even at very high (750 mg) doses^15^.

In addition to non-targeted GITR agonists, efforts are being made to develop tumor-targeted GITRL fusion proteins^11,16^, since there is evidence that the local delivery of GITR agonists can improve the therapeutic activity of GITR agonists^17^. Our group has worked for the last two decades on the antibody-based delivery of cytokine payloads to tumors and has characterized more than 100 fusion proteins until now^18^, but had never worked before with GITRL payloads. The F8 antibody, specific to the alternatively-spliced EDA domain of fibronectin^19^, is an attractive vehicle for pharmacodelivery applications. EDA is virtually absent from the adult human body (exception made for the female reproductive tract^20^) but represents an abundant component of the extracellular matrix of tumor-associated blood vessels^20–23^.

The targeted delivery of antibody-cytokine fusions, especially those featuring ligands of the TNF superfamily, to tumors has been demonstrated to be challenging in a number of cases^24^. Interaction of the cytokine moiety with cognate receptors outside the tumor can prevent accumulation of the antibody-cytokine fusion in the tumor^25^. While activation of the cytokine receptors in the periphery can lead to off-tumor toxicity, some anti-tumor activity might still be achieved^26^. In addition, many therapeutic proteins feature *N*-linked glycans which can be recognized by various glycoprotein receptors leading to degradation of the protein. For instance, terminal mannose or *N*-acetylglucosamine can be recognized by the mannose receptor expressed on macrophages and dendritic cells^27^. In addition, non-sialylated proteins with terminally exposed galactose moieties are recognized by hepatocytes expressing the asialoglycoprotein receptor^28^. Culture conditions^29,30^ during protein expression as well as the transfection method^31^ have a significant impact on the structure of the glycan making it difficult to obtain uniform glycosylation patterns across batches. Engineered aglycosylated protein variants that lack the consensus sequence for *N*-linked glycosylation offer a possibility to circumvent the problem of glycosylation-dependent protein degradation.

Here we report the development of an antibody-cytokine fusion featuring GITRL as a payload linked to the F8 antibody. Initial studies with the fusion protein featuring wild-type GITRL showed rapid degradation and lack of tumor accumulation *in vivo* which could be prevented by enzymatic deglycosylation of the protein. Therefore, fusion proteins featuring aglycosylated GITRL were developed by site-specific mutagenesis of *N*-linked glycosylation consensus sequence in the GITRL moiety. Similar to the enzymatically deglycosylated protein, the fusion protein featuring aglycosylated GITRL exhibited superior tumor-targeting properties compared to the fusion proteins featuring wild-type GITRL. Unfortunately, despite the improved targeting properties, no therapeutic anti-tumor activity could be observed.

## 2. MATERIALS AND METHODS

### 2.1 Cell lines

The murine cytotoxic T cell line CTLL-2 (ATCC^®^ TIB-214), the murine F9 teratocarcinoma cell line (ATCC^®^ CRL-1720) and the murine CT26 colon carcinoma cell line (ATCC^®^ CRL-2638) were obtained from ATCC, expanded and stored as cryopreserved aliquots in liquid nitrogen. The CTLL-2 cells were grown in RPMI-1640 (Gibco, #21875034) supplemented with 10% FBS (Gibco, #10270106), 1 X antibiotic-antimycoticum (Gibco, #15240062), 2 mM ultraglutamine (Lonza, #BE17-605E/U1), 25 mM HEPES (Gibco, #15630080), 50 μM β-mercaptoethanol (Sigma Aldrich) and 60 U/mL human IL-2 (Proleukin, Roche Diagnostics). The F9 teratocarcinoma cells were grown in DMEM (Gibco, high glucose, pyruvate, #41966-029) supplemented with 10% FBS (Gibco, #10270106) and 1 X antibiotic-antimycoticum (Gibco, #15240062) in flasks coated with 0.1% gelatin (Type B from Bovine Skin, Sigma Aldrich, #G1393). The CT26 colon carcinoma were grown in RPMI 1640 (Gibco, #21875034) supplemented with 10% FBS (Gibco, #10270106) and 1 X antibiotic-antimycoticum (Gibco, #15240062). The cells were passaged at the recommended ratios and never kept in culture for more than one month.

### 2.2 Mouse studies

Eight weeks old female 129/Sv and Balb/c mice were obtained from Janvier. The mice were kept in individually ventilated cages in groups of 5 mice per cage in a specific pathogen free facility. They received food and water ad libitum and the cages were changed once per week by trained caretakers. After at least one week of acclimatization, 7 – 10 x 10^6^ F9 cells (129/Sv) or 3 x 10^6^ CT26 cells (Balb/c) were subcutaneously implanted into the right flank. The tumor size was monitored daily by caliper measurements and the volume was calculated using the formula [length x width x width x 0.5]. The animals were euthanized when the tumor diameter exceeded 15 mm or when the tumor started to ulcerate. The animal experiments were carried out under the project license ZH04/2018 granted by the Veterinäramt des Kantons Zürich, Switzerland, in compliance with the Swiss Animal Protection Act (TSchG) and the Swiss Animal Protection Ordinance (TSchV).

### 2.3 Cloning

For PCR amplification of genetic sequences, the Phusion^®^ High-Fidelity PCR Master Mix (NewEnglandBiolabs, #M0532S) was used with a primer concentration of 200 nM. Genetic sequences encoding a single-chain trimer of the extracellular domain of murine GITRL (amino acids 46 – 170) linked by a short polypeptide linker (GGGSGGG) were obtained from Eurofins Genomics and introduced into a vector encoding the F8 or KSF antibody in the diabody format by Gibson Isothermal Assembly (NewEnglandBiolabs, NEBuilder^®^ HiFi DNA Assembly Master Mix, #E2621S). IgG fusions were cloned by fusing the sequence encoding GITRL to the genetic sequence encoding a chain of the IgG by PCR and introduced into the vector by restriction cloning. The genetic sequence encoding the diabody was replaced by the genetic sequence encoding the single-chain Fragment variable (scFv) by Gibson Isothermal Assembly and two domains of GITRL were removed by PCR followed by blunt-end ligation. Site-directed mutagenesis of the glycosylation consensus sequences were performed by Multichange Isothermal Mutagenesis (MISO) as described by Mitchell *et al*^32^. Brief, primers were designed including the desired point mutations, an upstream overhang of 14 bp and a downstream primer binding sequence with a melting point of around 60°C (calculated using tmcalculator.neb.com, Phusion^®^ High-Fidelity PCR Master Mix, 200 nM primer concentration). The fragments between two adjacent point mutations were amplified by PCR. Adjoining fragments were assembled by PCR and introduced into the backbone by Gibson Isothermal Assembly. Quality control of the plasmids was performed by Sanger Sequencing by Microsynth AG. The sequences of the proteins are provided in [**Supplementary Table 1**].

### 2.4 Protein production

Proteins were produced by transient transfection of CHO-S cells and purified by protein A affinity chromatography as described previously^33–35^. Brief, CHO-S cells were resuspended at a density of 4 mio cells/mL in ProCHO 4 medium (Lonza, #LZ-BE12-029Q) supplemented with 8 mM ultraglutamine (Lonza, #BE17-605E/U1), 1 X HT supplement (Gibco, # 41065012) and 1 X antibiotic-antimycoticum (Gibco, #15240062). DNA was added at a final concentration of 0.675 μg/mio cells and polyethylenimine (Polysciences, #23966-1) was added to a final concentration of 0.01 mg/mL. The cells were incubated in a shaking incubator at 31°C for 6 days before the supernatant was harvested and the proteins were purified by protein A affinity chromatography. The supernatant was filtered and applied to a protein A column at a speed of 2 mL/min at 4°C. The protein was washed by 100 – 200 mL of wash buffer A (100 mM NaCl, 0.5 mM EDTA, 0.1% Tween in PBS) and wash buffer B (500mM NaCl, 0.5 mM EDTA in PBS). The proteins were eluted in 10 mL of either 0.1 M glycine at pH 3 or 0.1 M Triethylamine, depending on the isoelectric point of the protein. The purified proteins were dialysed overnight against 3 L phosphate buffered saline (PBS). Quality control of the purified products included SDS-PAGE [**Supplementary Figure 1**], liquid chromatography-mass spectrometry (LC-MS) and size exclusion chromatography (SEC). Size exclusion chromatography was performed using an Äkta Pure FPLC system (GE Healthcare) with a Superdex S200 10/300 increase column at a flow rate of 0.75 mL/min (GE Healthcare) in PBS.

### 2.5 Binding measurements by Surface Plasmon Resonance

To evaluate the binding kinetics of the F8 moiety to EDA, a CM5 sensor chip (GE Healthcare) was coated with 500 resonance units of an EDA-containing recombinant fragment of fibronectin. The measurements were carried out with a Biacore S200 (GE Healthcare) setting the contact time to 3 min followed by a dissociation for 10 min and a regeneration of the chip using 10 mM HCl at a flow rate to 20 μL/min.

### 2.6 Binding measurements by Flow Cytometry

In order to assess the binding of the GITR moiety to cells expressing GITR, CTLL-2 cells were incubated with varying concentrations of the fusion proteins for 1 h. The bound protein was detected by addition of an excess of AlexaFluor488-labelled protein A (Thermofisher, #P11047) and subsequent measurement of the fluorescence using a Cytoflex Flow Cytometer. The mean fluorescence was normalized by subtracting the lowest measurement and dividing by the highest measurement. The resulting binding curve was fitted using the [Agonist] vs. response (three parameters) fit of the GraphPad Prism 7.0 a software to estimate the apparent K_D_.

### 2.7 NF-κB response assay

The development of the CTLL-2 reporter cell line was described previously^36^. Briefy, CTLL-2 reporter cells were washed with prewarmed HBSS (Gibco, #14175095) and grown for 6 – 9 h in growth medium without IL-2 prior to use in order to reduce the background signal. Cells were seeded in 96-well plates (50,000 cells/well) and growth medium containing varying concentrations of the antibody-GITRL conjugate was added. The cells were incubated at 37°C, 5% CO_2_ overnight. To assess luciferase production, 20 μL of the supernatant was transferred to an opaque 96-well plate (PerkinElmer, Optiplate-96, white, #6005290) and 80 μL 1 μg/mL Coelenterazine (Carl Roth AG, #4094.3) in phosphate buffered saline (PBS) was added. Luminescence at 466 nm was measured immediately. The relative luminescence was calculated by dividing the obtained results by the results obtained when no inducer was added. The resulting curve was fitted using the [Agonist] vs. response (three parameters) fit of the GraphPad Prism 7.0 a software to estimate the EC_50_.

### 2.8 Serum stability assay

In order to assess the *in vitro* stability of the proteins in mouse serum, the fusion proteins were diluted in mouse serum (Invitrogen) to 200 nM and incubated at 37°C for up to 48 h. After the incubation in mouse serum, a 10-fold dilution series was prepared and the above-described bioactivity assay was performed. The EC_50_ was estimated using the [Agonist] vs. response (three parameters) fit of the GraphPad Prism 7.0 a software.

### 2.9 Quantitative biodistribution

Quantitative biodistribution experiments were carried out as described previously19. Brief, F9 teratocarcinoma cells were cultivated and implanted into 129/Sv mice as described above. The fusion proteins were radioactively labelled with ^125^I using Chloramine-T. When the tumors reached a volume of 100 – 300 mm^3^ the mice were randomly assigned into groups of 3 mice and 10 – 15 μg of radioiodinated protein was injected into the lateral tail vein. The mice were sacrificed 24 h after the injection of the radiolabeled protein. The radioactivity of the excised organs was measured (Packard Cobra II Gamma Counter) and expressed as percentage of the injected dose per gram of tissue (%ID/g ± SD, *n* = 3). Enzymatic deglycosylation was performed overnight at 37°C using 3 U/μg protein PNGase F (NewEnglandBiolabs, #P0705S). As a negative control, equivalent antibody-cytokine fusions were used featuring the KSF antibody targeting hen egg lysozyme^22^[**Supplementary Figure 2**].

### 2.10 *Ex vivo* detection of fluorescently labelled proteins

Proteins were fluorescently labelled in a 0.1 M sodium carbonate buffer at pH 9.1 in the presence of excess Fluorescein Isothiocyanate overnight at 4°C. Uncojugated FITC was separated from the labelled proteins using PD-10 spin columns (Sigma Aldrich, PD Spintrap™ G25, #GE28-9180-04) according to the manufacturer’s recommendations. Approximately 100 μg of fluorescently labelled protein in PBS was injected into the lateral tail vein of tumor-bearing mice. The mice were sacrificed 24 h after the injection, the organs were embedded in NEG-50 cryoembedding medium (ThermoFisher, Richard-Allan-Scientific, #6502) and frozen. Of all samples, 8 μm cryosections were prepared and fixed in ice-cold acetone. The fixed slices were incubated with goat-anti-mouse CD31 (R&D system, #AF3628, 1:200) and rabbit-anti-FITC (Biorad, #4510-7804) followed by donkey-anti-goat-AF594 (Invitrogen, #A11058) and donkey-anti-rabbit-AF488 (Invitrogen, #A21206). Images were acquired using a Zeiss Axioscope 2 mot plus with an Axiocam 503 camera at a 200 X magnification in the RGB mode. The images were processed using the software ImageJ v1.52k.

### 2.11 Therapy studies

CT26 colon carcinoma cells were cultivated and implanted into Balb/c mice as described above. The tumor volume and the weight of the mice was monitored daily. When the tumor volume reached a value of 80 – 100 mm^3^ the mice were randomly assigned into groups of 5 animals in order to obtain uniform average tumor volumes for each group. Mice from each group were randomly distributed over the different cages. The mice received three cycles of intravenous injections of 200 μL of therapeutic agent. The experiment was performed under blinding conditions meaning that a second person prepared and labeled the therapeutic doses with a code which was only revealed to the researcher performing the animal experiments after the termination of the study. The control group received injections of phosphate buffered saline (PBS, Gibco, #1010023) only, one group received 200 μg of F8-GITRL followed by saline, one group received 200 μg of PD-1 inhibitor (BioXCell, clone 29F.1A12) followed by saline and the combination treatment group received 200 μg of PD-1 inhibitor followed by 200 μg of F8-GITRL [**Supplementary Table 4**].

### 2.12 Flow cytometry analysis of tumor infiltrating lymphocytes

CT26 tumor-bearing mice were treated as described above for the therapy [**Supplementary Table 4**]. The mice were sacrificed 24 h after the last therapeutic cycle and the tumor and draining lymph nodes were excised. The lymph nodes were mechanically disrupted on a 70 μm cell strainer. The tumors were cut into small pieces using a pair of surgical scissors and afterwards incubated in 5 mL of a digestion mix (RPMI-1640, 1 mg/mL collagenase II, 100 μg/mL DNaseI) in a shaking incubator at 37°C for 30 min. After the digestion of the extracellular matrix the tumor cells were passed through a cell strainer. The cells were harvested by centrifugation and red blood cells were removed using a red blood cell lysis buffer (Roche). Samples for intracellular staining were stained with zombie red for 15 min at room temperature. Surface staining was performed for 30 min on ice. For surface staining the following antibodies were used: αCD3-APC/Cy7 (Biolegend, #100222), αCD4-APC (Biolegend, #100412), αCD8-FITC (Biolegend, #100706), αCD8-APC/Cy7 (Biolegend, #100714), αNK1.1-PE (Biolegend, #108708), αCD62L-BV421 (Biolegend, #104436), αCD44-APC/Cy7 (Biolegend, #103028), αMHCII(IA/IE)-BV421 (Biolegend, #107631), αPD-1-BV421 (Biolegend, #109121), αCD39-APC (Biolegend, #143809), αCD127-APC (Biolegend, #135011), αCD226 (Biolegend, #128809), αTIGIT (Biolegend, #142111) and αGITR (Biolegend, #120205). The staining panel is depicted in [**Supplementary Table 5**]. The cells were washed twice in FACS buffer (0.5% BSA, 2 mM EDA, PBS). Samples that were stained for cell surface staining only were incubated for 5 min on ice in 7-AAD (Biolegend, #420404). Samples that were stained for intracellular markers were fixed and permeabilized using the eBioscience™ FoxP3/Transcription Factor Staining Buffer Set (Thermofisher, #00-5523-00) according to the manufacturer’s instructions. The samples were analyzed using a Beckmann Coulter Cytoflex and later processed using FlowJo v10.7.1. Statistical analysis was done using a regular two-way ANOVA followed by a Tukey’s multiple comparison test in GraphPad Prism v7. The gating strategy is shown in [**Supplementary Figure 6**]

## 3. RESULTS

### 3.1 Engineering of F8-GITRL fusion proteins

Five fusion proteins featuring wild-type murine GITRL linked to the F8 antibody were cloned and expressed. The different formats are schematically depicted in [**Figure 1 a**]. GITRL was fused to the F8 antibody both as a monomer and as a single-chain trimer, since forced trimerization has previously reported to increase the bioactivity of fusion proteins featuring members of the TNF superfamily^11^. The protein sequences are listed in [**Supplementary Table 1**]. The fusion proteins featuring wild-type GITRL as a payload were produced at yields ranging from 13 – 22 mg/L [**Supplementary Table 2**]. All variants yielded homogenous size exclusion chromatography profiles [**Figure 1 b**] and bound both to recombinant EDA (as measured by surface plasmon resonance) [**Figure 1 c**] and to the GITR-expressing murine cell line CTLL-2 (as measured by flow cytometry) [**Figure 1 d**]. In addition, measurements of the biological activity using an NF-κB reporter cell line showed that all formats triggered signal transduction in target cells [**Figure 1 e**]. Forced trimerization of the GITRL seemed to enhance the binding affinity and increased the biological activity in most cases [**Supplementary Table 3**]. Due to its favorable *in vitro* properties Format **1** was chosen for *in vivo* evaluation.

**Figure 1:**
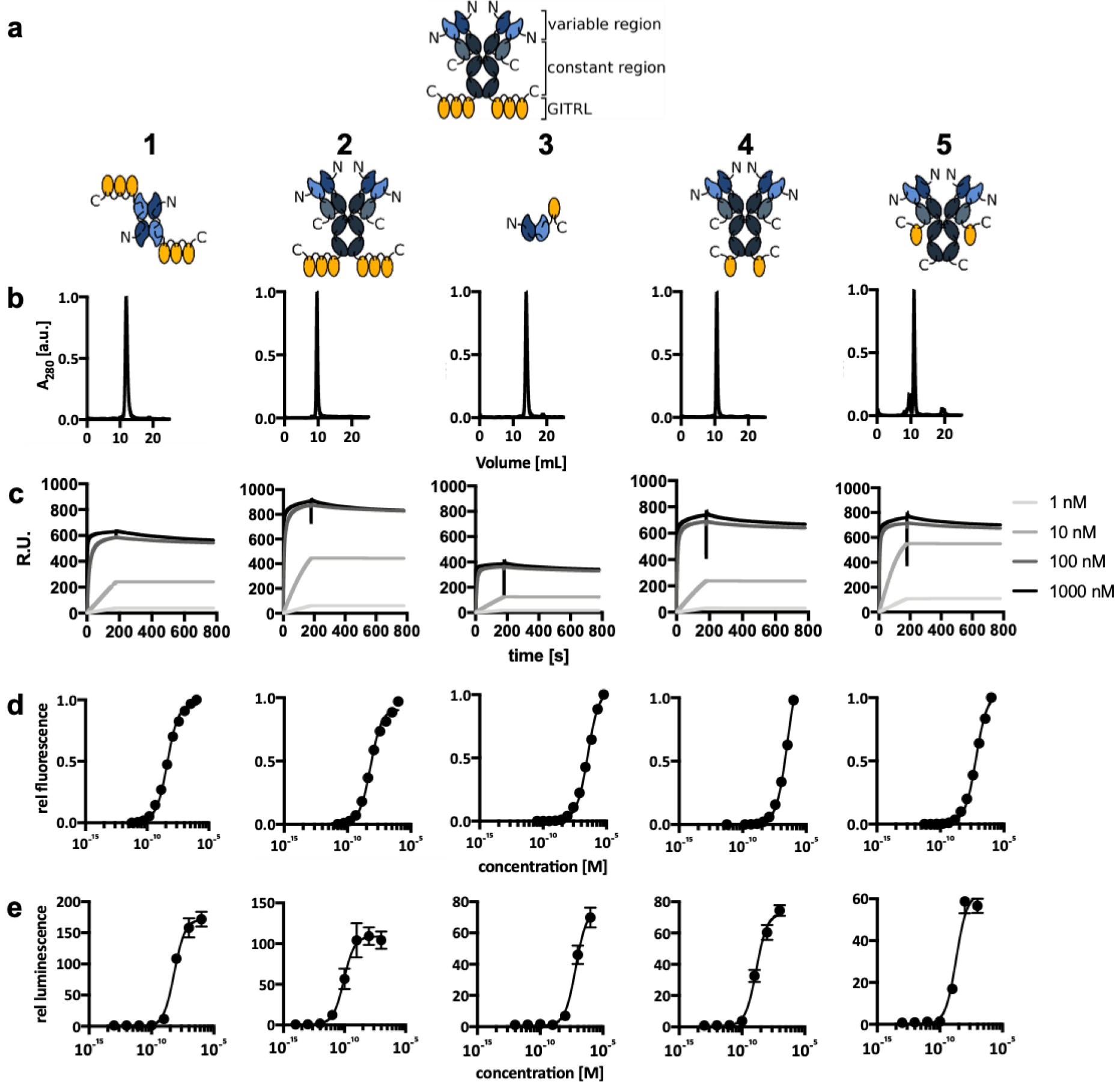
Screening of F8 antibody-cytokine fusion proteins featuring wild-type GITRL as immunomodulatory payload (a) Different formats of the F8 antibody such as diabody (Format **1**), full IgG (Formats **2** and **4**) or single-chain Fragment variable (scFv, Format **3**) were fused to GITRL either as a single-chain trimer (Formats **1** and **2**) or a monomer (Formats **3, 4** and **5**) (b) the size exclusion profile for each variant was measured using a Superdex S200 10/300 increase column (c) binding to recombinant EDA was measured by surface plasmon resonance (d) binding to GITRL was measured by flow cytometry on CTLL-2 cells (e) the *in vitro* bioactivity was measured using a CTLL-2 reporter cell line that secretes luciferase in response to NF-κB activation via GITR agonism. Data represents mean ± SD.

### 3.2 Quantitative biodistribution of F8-GITRL

A quantitative biodistribution of Format **1** revealed that the protein was rapidly cleared from the circulation without preferential tumor accumulation 24 h after the injection of the radioiodinated protein [**Figure 2**]. The tumor-targeting could be improved by enzymatic deglycosylation of the protein indicating that trapping of the antibody-cytokine fusion via the *N*-linked glycan lead to rapid degradation of the protein *in vivo*. In addition, after enzymatic deglycosylation also a higher %ID/g of 3% was measured in blood as compared to the native form for which only 0.1% ID/g were measured 24 h after administration of the protein. However, the dose at the tumor site varied considerably from mouse to mouse with one mouse reaching 5.6 %ID/g in the tumor and the other two reaching only 1-2 %ID/g of tumor.

**Figure 2:**
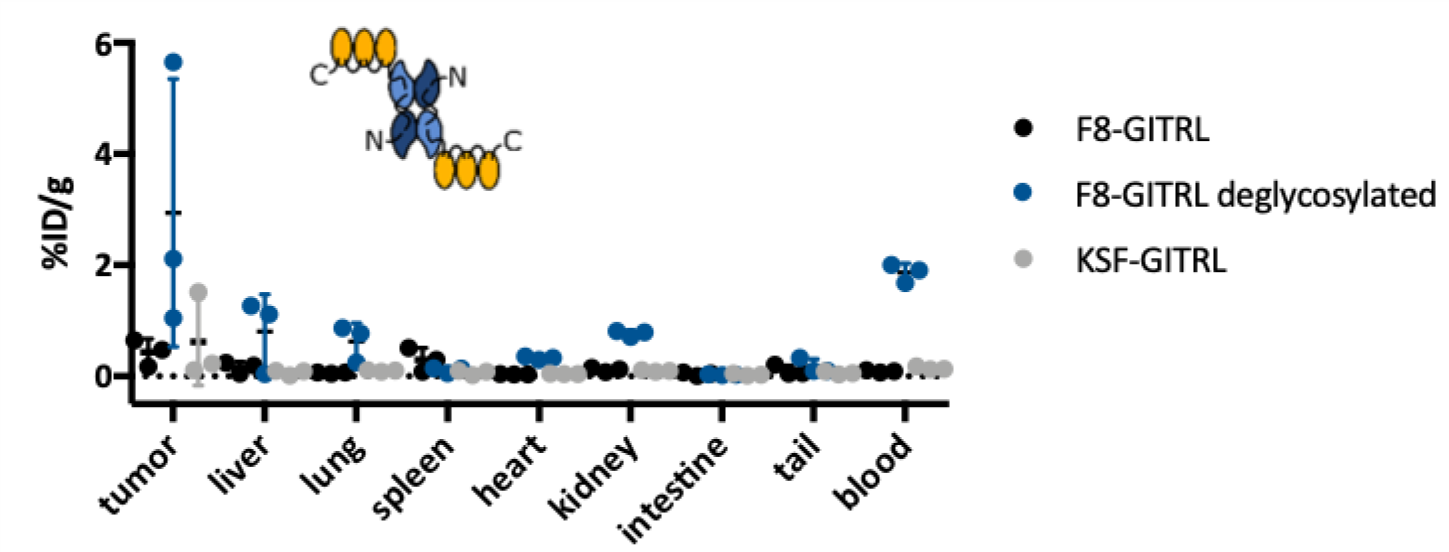
Quantitative Biodistribution of F8-GITRL in Format 1: The mice were sacrificed 24 h after intravenous administration of radiolabeled protein preparations and the radioactivity in the different organs was measured and expressed as percent injected dose per gram (%ID/g). The F8-GITRL protein was administered both in the native form and after enzymatic deglycosylation. As a negative control, a fusion protein featuring the KSF antibody which binds to hen egg lysozyme was used^22^. Data represents individual mice and mean ± SD (n = 3).

### 3.3 Engineering aglycosylated GITRL

In order to circumvent the need for enzymatic deglycosylation of F8-GITRL for *in vivo* applications, attempts were made to engineer aglycosylated variants. Therefore, the asparagine residues in the consensus sequence for *N*-linked glycosylation were sequentially removed by site-directed mutagenesis. At least in the absence of the glycan at position N74 of GITRL, the position N157 was occupied by a glycan and vice versa. Different combinations of double mutants were tested of which only the N74S and N157T could be expressed and was free of *N*-linked glycans [**Figure 3 a**]. However, the expression yield dropped from 13 mg/L for the fusion protein in Format **1** featuring wild-type GITRL to 1.5 mg/L for the aglycosylated variant [**Supplementary Table 2**]. More formats of antibody-cytokine fusions featuring the aglycosylated GITRL_N74S_N157T mutant (GITRL_mut) were developed and tested *in vitro* [**Figure 3 b, Supplementary Figure 3, Supplementary Figure 4**]. While Format **1^mut^** yielded equivalent results in terms of purity to the variants featuring wild-type GITRL, Format **2^mut^** was highly prone to aggregation and degradation [**Supplementary Figure 3**] and therefore not used for further studies. Format **6^mut^** exhibited substantial batch to batch variability in terms of protein quality. While some batches were of high quality, other batches contained substantial amounts of heavily degraded protein. In all cases, a significant decrease in expression yield was observed compared to the variant featuring wild-type GITRL [**Supplementary Table 2**]. By contrast, both the fusion protein in Format **1** featuring wild-type glycosylated GITRL and the one featuring aglycosylated GITRL retained the biological activity after incubation in mouse serum for up to 48 h at 37°C [**Supplementary Figure 5**].

**Figure 3:**
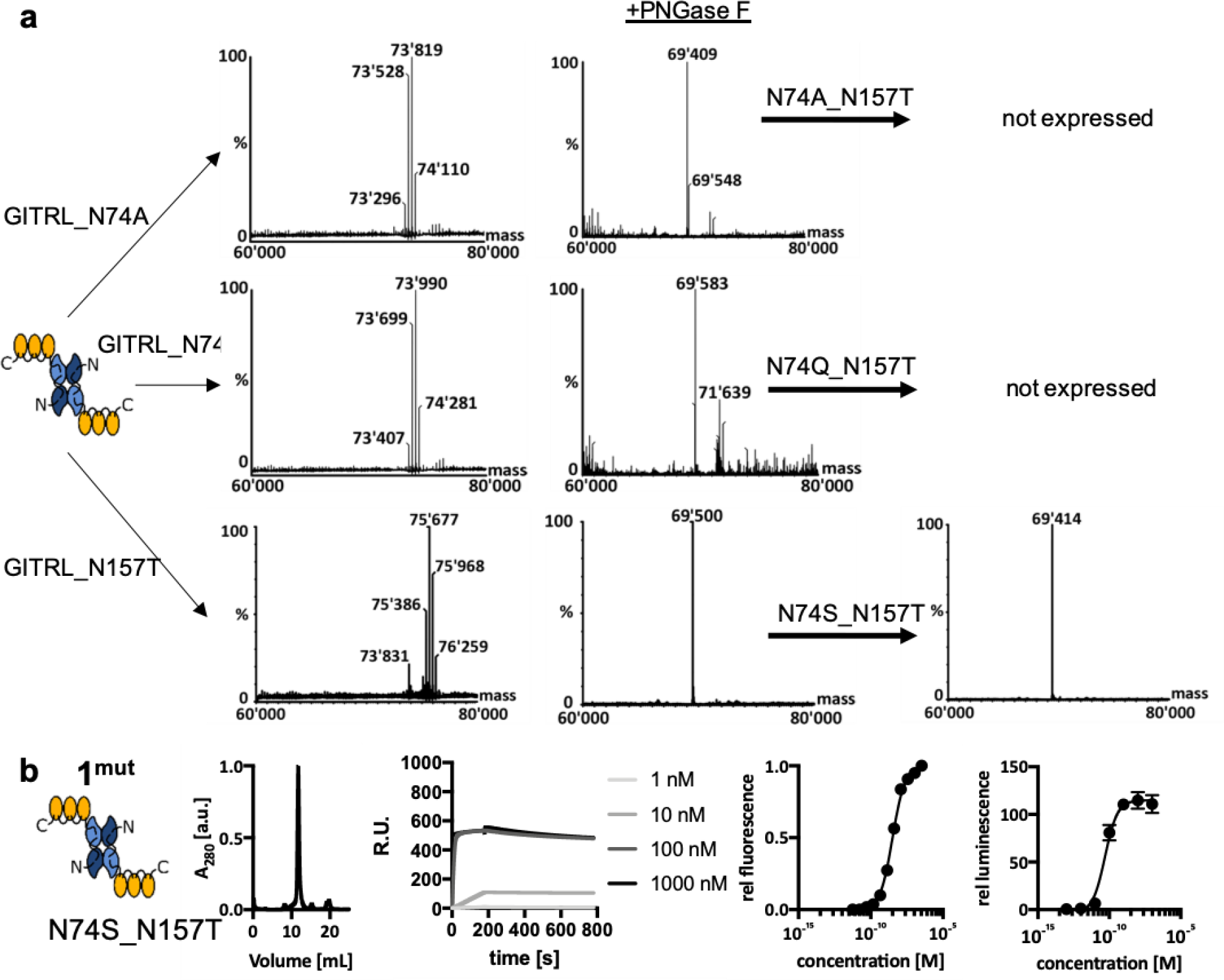
Development of aglycosylated variants of F8-GITRL (a) the asparagine residues N74 and N157 of GITRL were sequentially mutated and the resulting proteins were tested for the absence of *N*-linked glycans by mass spectrometry. Remaining *N*-linked glycans were removed by enzymatic deglycosylation with PNGase F. The single-site mutants retained a glycan while the double mutant N74S, N157T was the only non-glycosylated variant that could be expressed. (b) Characterization of the F8-GITRL fusion protein in Format 1^mut^ featuring aglycosylated GITRL_N74S_N157T

### 3.4 Biodistribution studies of aglycosylated F8-GITRL

Quantitative biodistribution studies with Format **1^mut^** featuring aglycosylated GITRL_N74S_N157T as payload showed that the fusion protein exhibited favorable tumor-targeting properties similar to what was observed after enzymatic deglycosylation of the fusion protein featuring wild-type GITRL [**Figure 4 a**]. As in the study in which enzymatically deglycosylated GITRL was used, a relatively high mouse to mouse variability in tumor uptake was observed. Selective accumulation of F8-GITRL featuring aglycosylated GITRL in the tumor was observed in F9 teratocarcinoma- and CT26 colon carcinoma-bearing mice 24 h after the injection of FITC-labelled protein [**Figure 4 b**]. Therefore, Format **1^mut^** featuring aglycosylated GITRL was selected for a therapy experiment in CT26 colon carcinoma-bearing mice.

**Figure 4:**
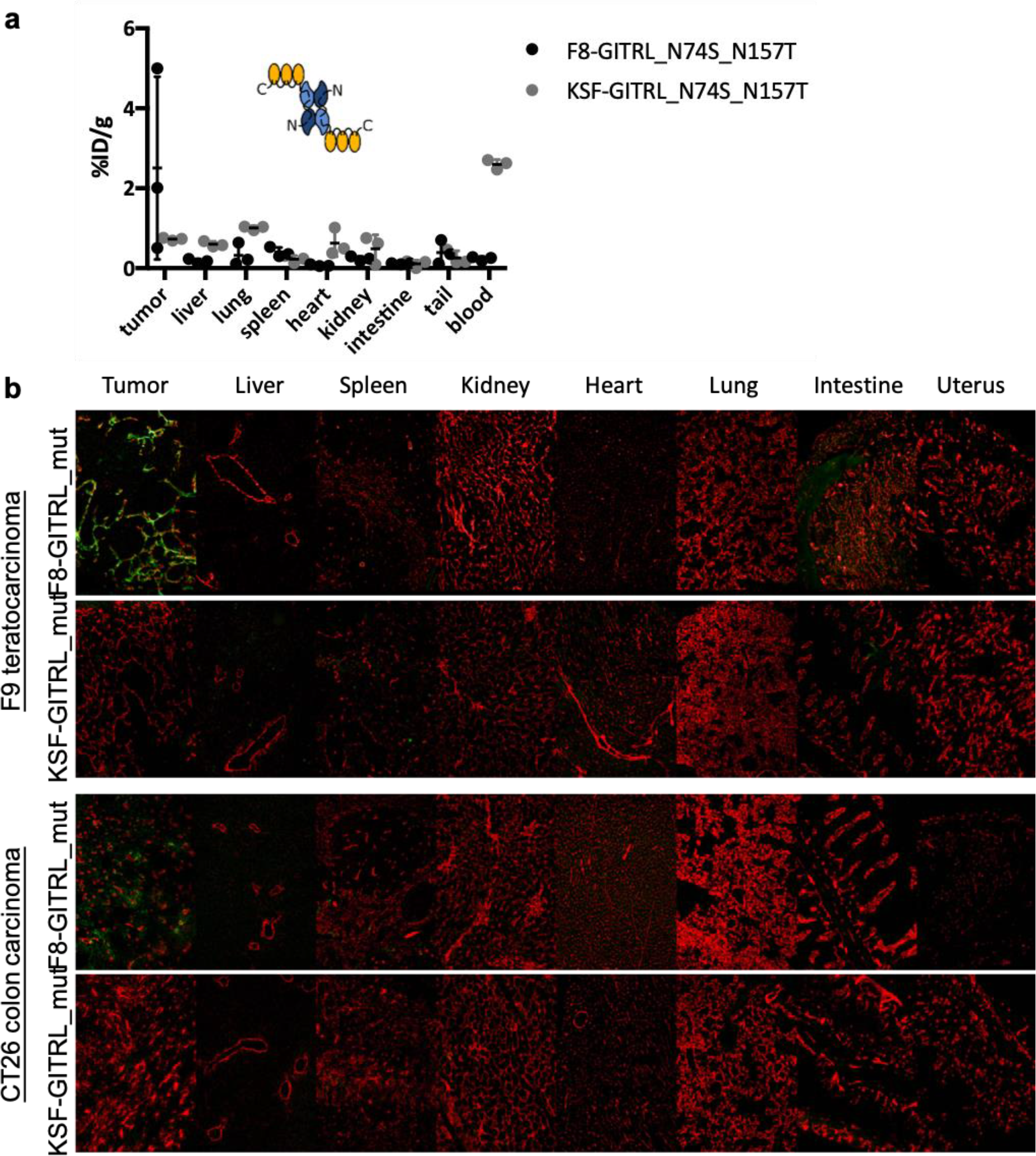
*In vivo* biodistribution of the F8 fusion protein in Format 1^mut^ featuring aglycosylated GITRL_N74S_N157T (GITRL_mut) (a) Mice were sacrificed 24 h after the intravenous administration of radiolabeled proteins and the radioactivity of the different organs was measured and expressed as % injected dose per gram of tissue (%ID/g). GITRL_N74S_NI57T (GITRL_mut) fused to the KSF antibody was used as negative control. The data represents individual measurements and mean ± SD. (b) Mice were sacrificed 24 h after intravenous administration of FITC-labelled protein preparations. The FITC-labelled proteins were detected *ex vivo* on cryosections. Green: αFITC, red: αCD31

### 3.5 Therapy studies using aglycosylated GITRL

The *in vivo* antitumor activity of F8-GITRL in Format 1^mut^ featuring aglycosylated GITRL_N74S_N157T was evaluated in CT26 colon carcinoma-bearing mice alone and in combination with a PD-1 inhibitor. The treatment schedule is shown in [Supplementary Table 4]. The different treatments did not show any effect on the growth rate of the tumors when compared to the group treated with saline only [Figure 5 a]. In general, the treatment was well tolerated as indicated by the absence of weight loss during the therapy [Figure 5 b]. Analysis of the tumor-infiltrating lymphocytes indicated an increase in antigen-presenting cells both in the tumor and the tumor-draining lymph nodes in the group treated with PD-1 inhibitor alone and in combination with F8-GITRL compared to the group treated with F8-GITRL only [Figure 5 c]. In addition, we observed an increase in the proportion of effector T cells in the tumor-draining lymph nodes amongst the CD8+ T cells specific for the tumor-rejection antigen AH137. The increase was higher in the groups treated with saline and F8-GITRL only [Figure 5 d]. In all treatment groups, the tumor-infiltrating CD8+ T cells were strongly positive for the exhaustion markers PD-1 and CD39 and expressed the negative costimulatory receptor TIGIT [Figure 5 e, f]. In addition, there was a non-significant tendency towards a higher proportion of regulatory T cells in the tumor-draining lymph nodes upon treatment with a PD-1 inhibitor [Figure 5 g]. In conclusion, the treatment failed to reinvigorate the anti-tumor immune response.

**Figure 5:**
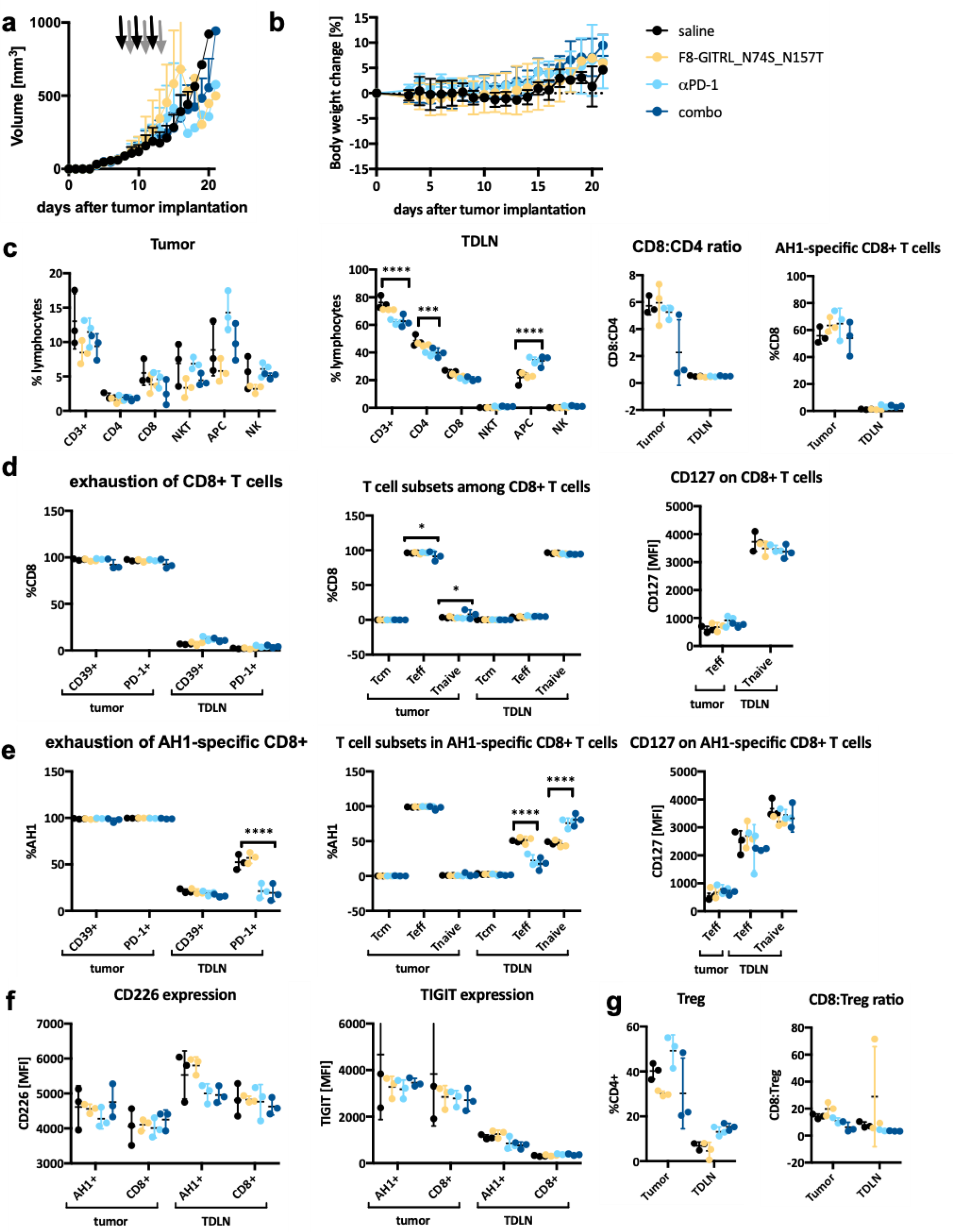
Therapy studies and immune infiltrate analysis in CT26 colon carcinoma-bearing mice (a) CT26 tumor-bearing mice received three cycles of injections of either twice saline, F8-GITRL_N74S_N157T followed by saline, αPD-1 followed by saline or αPD-1 followed by F8-GITRL_N74S_N157T. The black arrows indicate the first injection of each cycle whereas the grey arrows indicate the second injection of each cycle. The tumor volume is depicted as average +SD for each group (n= 5). (b) The average body weight change after tumor cell implantation is depicted as mean ±SD. (c) The composition of tumor-infiltrating lymphocytes and the tumor-draining lymph nodes (TDLN) (d) the phenotype of CD8+ T cells in the tumor and TDLN (e) the phenotype of CD8+ T cells specific for the tumor-rejection antigen AH1 in the tumor and TDLN (f) expression of CD226 and TIGIT on CD8+ T cells in the tumor and TLDN (g) regulatory T cells in the tumor and TDLN as analyzed by flow cytometry. Data represents individual measurements and mean ± SD (n = 3). Statistical analysis was performed by a regular two-way ANOVA with Tukey’s post-test in Graph Pad prism 7 (p < 0.0001: ****, p < 0.001: ***, p < 0.01: **, p < 0.05: *, p > 0.05: not significant)

## 4. DISCUSSION

The clinical success of immune checkpoint inhibitors in a subset of patients highlighted the need for additional immunomodulatory treatments in order to reach therapeutic success in a wider population of cancer patients. In the recent years, GITR has emerged as a promising target to deliver costimulatory signals to effector T cells and to rescue this subset from suppression by regulatory T cells^6^. Several lines of evidence showed potent synergistic antitumor activity of GITR agonists in combination with PD-1 inhibitors^10,38^. A number of GITR agonists are currently investigated in clinical trials.

In this study, we developed a fusion protein consisting of murine GITRL and the F8 antibody targeting the EDA-positive splice isoform of fibronectin which is a marker of the tumor neovasculature^19,21^. We demonstrated that the *in vivo* stability and tumor-targeting properties of the F8-GITRL fusion protein featuring wild-type murine GITRL as a payload were hampered by glycan-mediated clearance of the protein. Although glycan-mediated clearance could to some extent be prevented by enzymatic deglycosylation and site-specific mutagenesis of the *N*-glycosylation consensus sequence, a substantial mouse-to-mouse variability in tumor uptake was observed. In addition, the yield of fusion proteins featuring aglycosylated GITRL was significantly lower than the constructs featuring wild-type GITRL. Given the importance of *N*-linked glycosylation for folding and quality control of secreted proteins^39^, the lower yield could indicate a reduction in protein folding efficiency during the production in CHO cells. In addition, the removal of the *N*-linked glycan could unmask cleavage sites for serum proteases or render the protein more prone to denaturation. However, both the wild-type and the aglycosylated mutant of GITRL fused to F8 in the diabody format retained the *in vitro* biological activity when incubated in mouse serum for up to 48 h, indicating that the absence of a glycan did not significantly impair the stability of the fully folded protein. While this is a rather indirect readout for the stability of the protein, we speculated that upon degradation or unfolding, the protein would lose its bioactivity. In addition, a protein which does not retain *in vitro* bioactivity under the given conditions is also not expected to have any therapeutic activity *in vivo*.

Treatment with F8-GITRL, alone or in combination with a PD-1 inhibitor, failed to mediate an antitumor immune response in CT26 colon carcinoma-bearing mice. The presence of CD8+ T cells specific for the retroviral tumor rejection antigen AH1 among the tumor infiltrating lymphocytes was confirmed by flow cytometry [**Figure 5 c**]. Previous reports had shown that AH1-specific CD8+ T cells constitute the major drivers of the antitumor immune response against CT26 colon carcinoma^37,40,41^. In line with previous reports, tumor infiltrating CD8+T cells were positive for the exhaustion markers^42^ such as CD39 and PD-1 [**Figure 5 d, e**]. Previous studies had shown that the combination of GITR agonistic antibodies and PD-1 inhibitors could revert the dysfunctional state of tumor-specific CD8+ T cells by inducing the downregulation of TIGIT and the upregulation of the costimulatory receptor CD226 in murine models of cancer10. We did not observe any modulation of the expression of TIGIT or CD226 on tumor-infiltrating CD8+T cells. We also did not observe any modulation in the number of regulatory T cells or the ratio of effector T cells to regulatory T cells which was suggested as a biomarker for anti-GITR activity in other reports^38^.

The biological role of GITR as a target for anti-cancer intervention is questionable, also in light of the findings of our work. A number of preclinical studies has previously reported a promising anti-tumor activity of GITR agonists^6,43^. However, a Phase I clinical trial with a GITRL-Fc fusion (MEDI1873^44^) failed to demonstrate objective responses, in spite of the fact that a very broad dose range was tested (i.e., between 1.5 mg and 750 mg per injection)^15^. Similarly, a lack of clinical benefit was reported for a number of GITR agonistic antibodies such as TRX518^38^, AMG228^45^, BMS-986156^46^ and MK-1248^47^. Thus

In summary, we could identify a molecular format and suitable glycoengineering strategies, that allowed the creation of a novel fusion protein (F8-GITRL) with promising tumor-homing properties, as revealed by quantitative biodistribution analysis and by *ex vivo* immunofluorescence studies in tumor-bearing mice. However, the lack of anticancer activity of the fusion protein *in vivo* casts doubts about the potential of GITRL as a payload for tumor therapy strategies.

## ACKNOWLEDGEMENTS

Financial support by the ETH Zürich, the Swiss National Science Foundation (grant number 310030_182003/1), the European Research Council (ERC) under the European Union’s Horizon 2020 research and innovation program (grant agreement 670603), and the Federal Commission for Technology and Innovation (KTI, grant number 12803.1 VOUCH-LS) is gratefully acknowledged. The authors gratefully acknowledge Fiona Ammann and Sabrina Müller for their technical assistance.

## AUTHOR CONTRIBUTIONS

**Jacqueline Mock**: Conceptualization, Methodology, Project administration, Investigation, Writing – Original Draft. **Itzel Astiazaran Rascon:** Investigation. **Marco Stringhini:** Investigation, Methodology. **Marco Catalano:** Investigation. **Dario Neri**: Supervision, Funding acquisition, Writing – Original Draft, Resources, Conceptualization.

## DATA AVAILABILITY STATEMENT

The datasets generated during and/or analyzed during the current study are available from the corresponding author on reasonable request.

