## SupplementaryData for "Engineering murine GITRL for antibody-mediated delivery to tumor-associated blood vessels"

**F8(dDb)-(4-GITRL)$_3$**: F8_VH-*linker*-F8_VL-*linker*-GITRL-*linker*-GITRL-*linker*-GITRL (Format **1**)
EVQLLESGGGLVQPGGSLRLSCAASGFTFSLFTMSWVRQAPGKGLEWVSAISGSGGSTYYADSVKGRFTISRDNSKNTLYLQ
MNSLRAEDTAVYYCAKSTHLYLFDYWGQGTLVTVSS-*GGSGG*-EIVLTQSPGTLSLSPGERATLSCRASQSVSMPFLAWYQQ
KPGQAPRLLIYGASSRATGIPDRFSGSGSGTDFTLTISRLEPEDFAVYYCQQMRGRPPTFGQGTKVEIK-*SSSSGSSSSGSSSSG*
– PTAIESCMVKFELSSSKWHMTSPKPHCV**N**TTSDGKLKILQSGTYLIYGQVIPVDKKYIKDNAPFVVQIYKKNDVLQTLMND
FQILPIGGVYELHAGDNIYLKFNSKDHIQK**N**NTYWGIILMPDLP-*GGGSGGG*-PTAIESCMVKFELSSSKWHMTSPKPHCV**N**
TTSDGKLKILQSGTYLIYGQVIPVDKKYIKDNAPFVVQIYKKNDVLQTLMNDFQILPIGGVYELHAGDNIYLKFNSKDHIQK**N**N
TYWGIILMPDLP-*GGGSGGG*-PTAIESCMVKFELSSSKWHMTSPKPHCV**N**TTSDGKLKILQSGTYLIYGQVIPVDKKYIKDNA
PFVVQIYKKNDVLQTLMNDFQILPIGGVYELHAGDNIYLKFNSKDHIQK**N**NTYWGIILMPDLP
**F8(IgG)-(GITRL)$_3$_HC**: VL-CL** VH-CH-*linker*-GITRL-*linker*-GITRL-*linker*-GITRL (Format **2**)
EIVLTQSPGTLSLSPGERATLSCRASQSVSMPFLAWYQQKPGQAPRLLIYGASSRATGIPDRFSGSGSGTDFTLTISRLEPEDF
AVYYCQQMRGRPPTFGQGTKVEIKRTVAAPSVFIFPPSDEQLKSGTASVVCLLNNFYPREAKVQWKVDNALQSGNSQESV
TEQDSKDSTYSLSSTLTLSKADYEKHKVYACEVTHQGLSSPVTKSFNRGEC**
EVQLLESGGGLVQPGGSLRLSCAASGFTFSLFTMSWVRQAPGKGLEWVSAISGSGGSTYYADSVKGRFTISRDNSKNTLYLQ
MNSLRAEDTAVYYCAKSTHLYLFDYWGQGTLVTVSSASTKGPSVFPLAPSSKSTSGGTAALGCLVKDYFPEPVTVSWNSGAL
TSGVHTFPAVLQSSGLYSLSSVVTVPSSSLGTQTYICNVNHKPSNTKVDKKVEPKSCDKTHTCPPCPAPELLGGPSVFLFPPKP
KDTLMISRTPEVTCVVVDVSHEDPEVKFNWYVDGVEVHNAKTKPREEQYNSTYRVVSVLTVLHQDWLNGKEYKCKVSNKA
LPAPIEKTISKAKGQPREPQVYTLPPSRDELTKNQVSLTCLVKGFYPSDIAVEWESNGQPENNYKTTPPVLDSDGSFFLYSKLT
VDKSRWQQGNVFSCSVMHEALHNHYTQKSLSLSPGK-*SSSSGSSSSGSSSSG*-
PTAIESCMVKFELSSSKWHMTSPKPHCV**N**TTSDGKLKILQSGTYLIYGQVIPVDKKYIKDNAPFVVQIYKKNDVLQTLMNDF
QILPIGGVYELHAGDNIYLKFNSKDHIQK**N**NTYWGIILMPDLP-*GGGSGGG*-PTAIESCMVKFELSSSKWHMTSPKPHCV**N**T
TSDGKLKILQSGTYLIYGQVIPVDKKYIKDNAPFVVQIYKKNDVLQTLMNDFQILPIGGVYELHAGDNIYLKFNSKDHIQK**N**NT
YWGIILMPDLP-*GGGSGGG*-PTAIESCMVKFELSSSKWHMTSPKPHCV**N**TTSDGKLKILQSGTYLIYGQVIPVDKKYIKDNAP
FVVQIYKKNDVLQTLMNDFQILPIGGVYELHAGDNIYLKFNSKDHIQK**N**NTYWGIILMPDLP
**F8(scFv)-GITRL**: F8_VH-*linker*-F8_VL-*linker*-GITRL (Format **3**)
EVQLLESGGGLVQPGGSLRLSCAASGFTFSLFTMSWVRQAPGKGLEWVSAISGSGGSTYYADSVKGRFTISRDNSKNTLYLQ
MNSLRAEDTAVYYCAKSTHLYLFDYWGQGTLVTVSS-*GGGGSGGGGSGGGG*-EIVLTQSPGTLSLSPGERATLSCRASQSVS
MPFLAWYQQKPGQAPRLLIYGASSRATGIPDRFSGSGSGTDFTLTISRLEPEDFAVYYCQQMRGRPPTFGQGTKVEIK-*SSSS*
*GSSSSGSSSS*G- PTAIESCMVKFELSSSKWHMTSPKPHCV**N**TTSDGKLKILQSGTYLIYGQVIPVDKKYIKDNAPFVVQIYKKN
DVLQTLMNDFQILPIGGVYELHAGDNIYLKFNSKDHIQK**N**NTYWGIILMPDLP
**F8(IgG)-GITRL_HC**: VL-CL** VH-CH-*linker*-GITRL (Format **4**)
EIVLTQSPGTLSLSPGERATLSCRASQSVSMPFLAWYQQKPGQAPRLLIYGASSRATGIPDRFSGSGSGTDFTLTISRLEPEDF
AVYYCQQMRGRPPTFGQGTKVEIKRTVAAPSVFIFPPSDEQLKSGTASVVCLLNNFYPREAKVQWKVDNALQSGNSQESV
TEQDSKDSTYSLSSTLTLSKADYEKHKVYACEVTHQGLSSPVTKSFNRGEC**
EVQLLESGGGLVQPGGSLRLSCAASGFTFSLFTMSWVRQAPGKGLEWVSAISGSGGSTYYADSVKGRFTISRDNSKNTLYLQ
MNSLRAEDTAVYYCAKSTHLYLFDYWGQGTLVTVSSASTKGPSVFPLAPSSKSTSGGTAALGCLVKDYFPEPVTVSWNSGAL
TSGVHTFPAVLQSSGLYSLSSVVTVPSSSLGTQTYICNVNHKPSNTKVDKKVEPKSCDKTHTCPPCPAPELLGGPSVFLFPPKP
KDTLMISRTPEVTCVVVDVSHEDPEVKFNWYVDGVEVHNAKTKPREEQYNSTYRVVSVLTVLHQDWLNGKEYKCKVSNKA
LPAPIEKTISKAKGQPREPQVYTLPPSRDELTKNQVSLTCLVKGFYPSDIAVEWESNGQPENNYKTTPPVLDSDGSFFLYSKLT
VDKSRWQQGNVFSCSVMHEALHNHYTQKSLSLSPGK-*SSSSGSSSSGSSSSG*- PTAIESCMVKFELSSSKWHMTSPKPHCV
**N**TTSDGKLKILQSGTYLIYGQVIPVDKKYIKDNAPFVVQIYKKNDVLQTLMNDFQILPIGGVYELHAGDNIYLKFNSKDHIQK**N**
NTYWGIILMPDLP
**F8(IgG)-GITR_LC**: VL-CL-*linker*-GITRL** VH-CH (Format **5**)
EIVLTQSPGTLSLSPGERATLSCRASQSVSMPFLAWYQQKPGQAPRLLIYGASSRATGIPDRFSGSGSGTDFTLTISRLEPEDF
AVYYCQQMRGRPPTFGQGTKVEIKRTVAAPSVFIFPPSDEQLKSGTASVVCLLNNFYPREAKVQWKVDNALQSGNSQESV
TEQDSKDSTYSLSSTLTLSKADYEKHKVYACEVTHQGLSSPVTKSFNRGEC-*SSSSGSSSSGSSSSG*- PTAIESCMVKFELSSSK
WHMTSPKPHCV**N**TTSDGKLKILQSGTYLIYGQVIPVDKKYIKDNAPFVVQIYKKNDVLQTLMNDFQILPIGGVYELHAGDNI
YLKFNSKDHIQK**N**NTYWGIILMPDLP **
EVQLLESGGGLVQPGGSLRLSCAASGFTFSLFTMSWVRQAPGKGLEWVSAISGSGGSTYYADSVKGRFTISRDNSKNTLYLQ
MNSLRAEDTAVYYCAKSTHLYLFDYWGQGTLVTVSSASTKGPSVFPLAPSSKSTSGGTAALGCLVKDYFPEPVTVSWNSGAL
TSGVHTFPAVLQSSGLYSLSSVVTVPSSSLGTQTYICNVNHKPSNTKVDKKVEPKSCDKTHTCPPCPAPELLGGPSVFLFPPKP
KDTLMISRTPEVTCVVVDVSHEDPEVKFNWYVDGVEVHNAKTKPREEQYNSTYRVVSVLTVLHQDWLNGKEYKCKVSNKA
LPAPIEKTISKAKGQPREPQVYTLPPSRDELTKNQVSLTCLVKGFYPSDIAVEWESNGQPENNYKTTPPVLDSDGSFFLYSKLT
VDKSRWQQGNVFSCSVMHEALHNHYTQKSLSLSPGK
**KSF(dDb)-(4-GITRL)$_3$**: KSF_VH-*linker*-KSF_VL-*linker*-GITRL-*linker*-GITRL-*linker*-GITRL (Format **1**)

EVQLLESGGGLVQPGGSLRLSCAASGFTFSSYAMSWVRQAPGKGLEWVSAISGSGGSTYYADSVKGRFTISRDNSKNTLYL
QMNSLRAEDTAVYYCAKSPKVSLFDYWGQGTLVTVSS-*GGSGG*-SELTQDPAVSVALGQTVRITCQGDSLRSYYASWYQQ
KPGQAPVLVIYGKNNRPSGIPDRFSGSSSGNTASLTITGAQAEDEADYYCNSSPLNRLAVVFGGGTKLTVLG-*SSSSGSSSSGS*
*SSSG* – PTAIESCMVKFELSSSKWHMTSPKPHCV**N**TTSDGKLKILQSGTYLIYGQVIPVDKKYIKDNAPFVVQIYKKNDVLQTL
MND FQILPIGGVYELHAGDNIYLKFNSKDHIQK**N**NTYWGIILMPDLP-*GGGSGGG*-PTAIESCMVKFELSSSKWHMTSPKP
HCV**N**TTSDGKLKILQSGTYLIYGQVIPVDKKYIKDNAPFVVQIYKKNDVLQTLMNDFQILPIGGVYELHAGDNIYLKFNSKDHI
QK**N**NTYWGIILMPDLP-*GGGSGGG*-PTAIESCMVKFELSSSKWHMTSPKPHCV**N**TTSDGKLKILQSGTYLIYGQVIPVDKKY
IKDNAPFVVQIYKKNDVLQTLMNDFQILPIGGVYELHAGDNIYLKFNSKDHIQK**N**NTYWGIILMPDLP

Supplementary Table 1: Sequences of the fusion proteins that were developed in this study. The asparagine residues N74 and N157 of GITRL that were mutated to S and T are highlighted in bold. The consensus sequences for *N*-linked glycosylation are underlined.

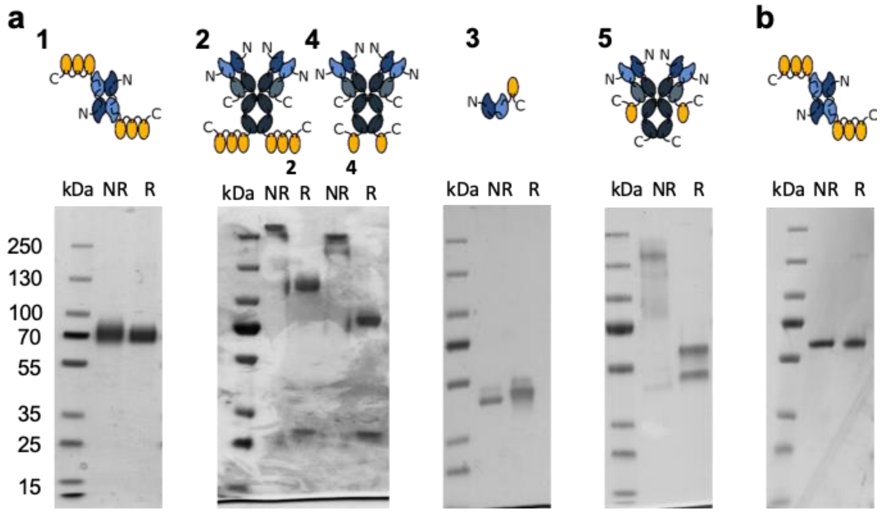


Supplementary Figure 1: SDS PAGE analysis of the F8-GITRL fusion proteins featuring (a) wild-type GITRL and (b) aglycosylated GITRL (NR: non-reducing sample buffer, R: reducing sample buffer)

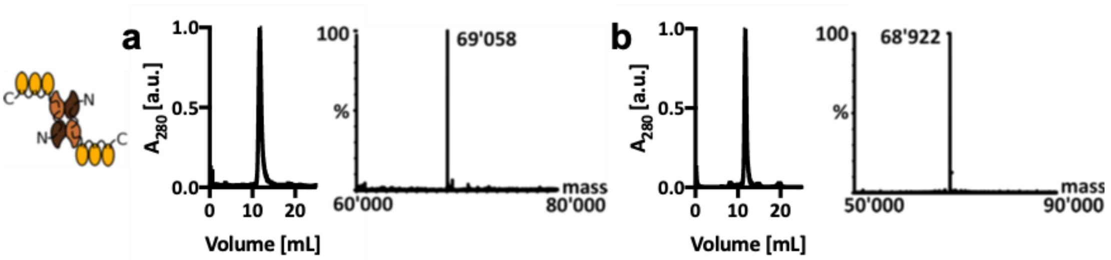


Supplementary Figure 2: Characterization of the KSF-GITRL fusion proteins that were used as negative control in the quantitative biodistribution studies (a) Size exclusion chromatogram and LC-MS spectrum of KSF-GITRL in Format 1 featuring wild-type GITRL (b) Size exclusion chromatogram and LC-MS spectrum of KSF-GITRL_N74S_N157T in Format $1^{mut}$ featuring the aglycosylated GITRL mutant

| Format | Yield WT-GITRL [mg/L] | Yield GITRL_N74S_N157T [mg/L] |
|---|---|---|
| 1: F8(dDb)-$(GITRL)_3$ | 13 | 1.5 |
| 2: F8(IgG)-$(GITRL)_3$_HC | 22 | n/a |
| 3: F8(scFv)-GITRL | 19 | 5 |
| 6: F8(IgG)-$(GITRL)_3$_LC | n/a | 4.7 |

Supplementary Table 2: Expression yields from the purification of the different F8-GITRL fusion proteins featuring wild-type and aglycosylated GITRL as payloads. The proteins were expressed in CHO cells as described in the methods section.

| Format | apparent $K_D$ [M] | | | $EC_{50}$ [M] | | |
|---|---|---|---|---|---|---|
| 1: F8(dDb)-$(GITRL)_3$ | 4.4E-09 | ± | 4.4E-10 | 4.7E-09 | ± | 4.8E-10 |
| 2: F8(IgG)-$(GITRL)_3$_HC | 6.7E-09 | ± | 8.1E-10 | 8.9E-11 | ± | 1.9E-11 |
| 3: F8(scFv)-GITRL | 3.4E-07 | ± | 3.1E-08 | 7.0E-08 | ± | 9.7E-09 |
| 4: F8(IgG)-GITRL_HC | 3.3E-07 | ± | 2.2E-08 | 5.8E-09 | ± | 9.2E-10 |
| 5: F8(IgG)-GITRL_LC | 6.4E-08 | ± | 5.6E-09 | 2.0E-09 | ± | 4.2E-10 |
| $1^{mut}$: F8(dDb)-$(GITRL\_N74S\_N157T)_3$ | 3.2E-09 | ± | 2.5E-10 | 5.0E-11 | ± | 8.6E-12 |
| $6^{mut}$: F8(IgG)-$(GITRL_N74S_N157T)_3$_LC | 1.5E-09 | ± | 3.6E-10 | 1.5E-11 | ± | 2.6E-12 |

Supplementary Table 3: Binding affinity and *in* vitro bioactivity of the F8-GITRL as measured by flow cytometry binding to CTLL-2 cells and using the CTLL-2 NF-κB reporter cell line.

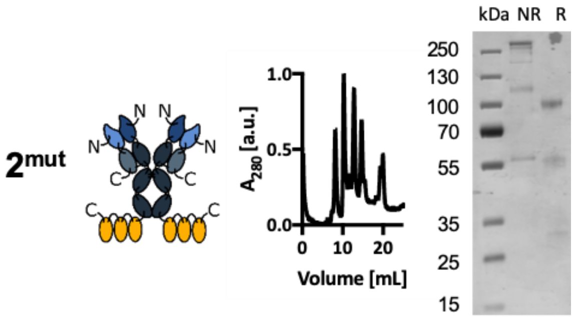


Supplementary Figure 3: The fusion protein consisting of aglycosylated GITRL_N74S_N157T fused to the heavy chain of the F8 antibody (Format $2^{mut}$) yielded a heavily degraded protein as can be seen on the size exclusion chromatogram and on SDS PAGE (NR: non-reducing sample buffer, R: reducing sample buffer).

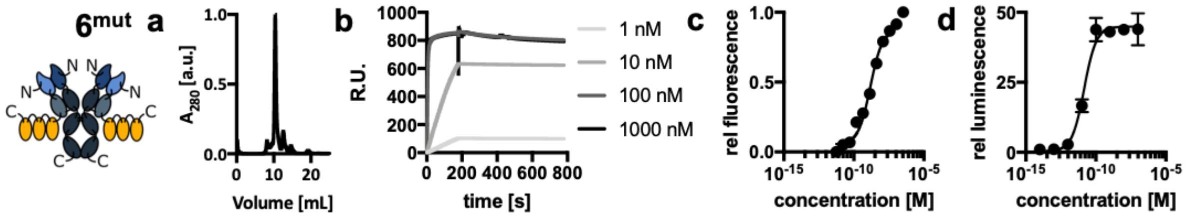


Supplementary Figure 4: Characterization of aglycosylated GITRL_N74S_N157T fused to the light chain of the F8 antibody in the IgG format (Format $6^{mut}$) (a) size exclusion chromatogram (b) surface plasmon resonance measurement of the binding to EDA (c) flow cytometry measurement of the binding to CTLL-2 (d) *in vitro* bioactivity

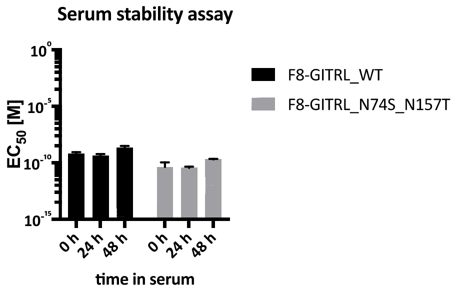


Supplementary Figure 5: *In vitro* serum stability assay, the protein was incubated in mouse serum at 37°C for up to 48 h and the biological activity was measured using the NF-κB reporter cell line.

| | Days after tumor implantation | | | | | |
|---|---|---|---|---|---|---|
| **group** | **8** | **9** | **10** | **11** | **12** | **13** |
| Saline | PBS | PBS | PBS | PBS | PBS | PBS |
| F8-GITRL | F8-GITRL | PBS | F8-GITRL | PBS | F8-GITRL | PBS |
| $\alpha$PD-1 | $\alpha$PD-1 | PBS | $\alpha$PD-1 | PBS | $\alpha$PD-1 | PBS |
| combo | $\alpha$PD-1 | F8-GITRL | $\alpha$PD-1 | F8-GITRL | $\alpha$PD-1 | F8-GITRL |

Supplementary Table 4: Treatment schedule for the therapy and infiltrate analysis of CT26 colon carcinoma-bearing mice using F8-GITRL_N74S_N157T in Format $1^{mut}$ and a PD-1 inhibitor

| | **FITC** | **APC/Cy7** | **PE** | **APC** | **BV421** | **Live/dead** |
|---|---|---|---|---|---|---|
| Stain 1 | CD8 | CD3 | NK1.1 | CD4 | IA/IE | 7-AAD |
| Stain 2 | CD8 | CD44 | AH1 | CD127 | CD62L | 7-AAD |
| Stain 3 | CD8 | - | AH1 | CD39 | PD-1 | 7-AAD |
| Stain 4 | CD8 | - | AH1 | CD226 | TIGIT | 7-AAD |
| Stain 5 | GITR | CD8 | - | CD4 | FoxP3 | Zombie red |

Supplementary Table 5: Design of the 5-color panel for the analysis of tumor infiltrating lymphocytes and tumor-draining lymph nodes by flow cytometry

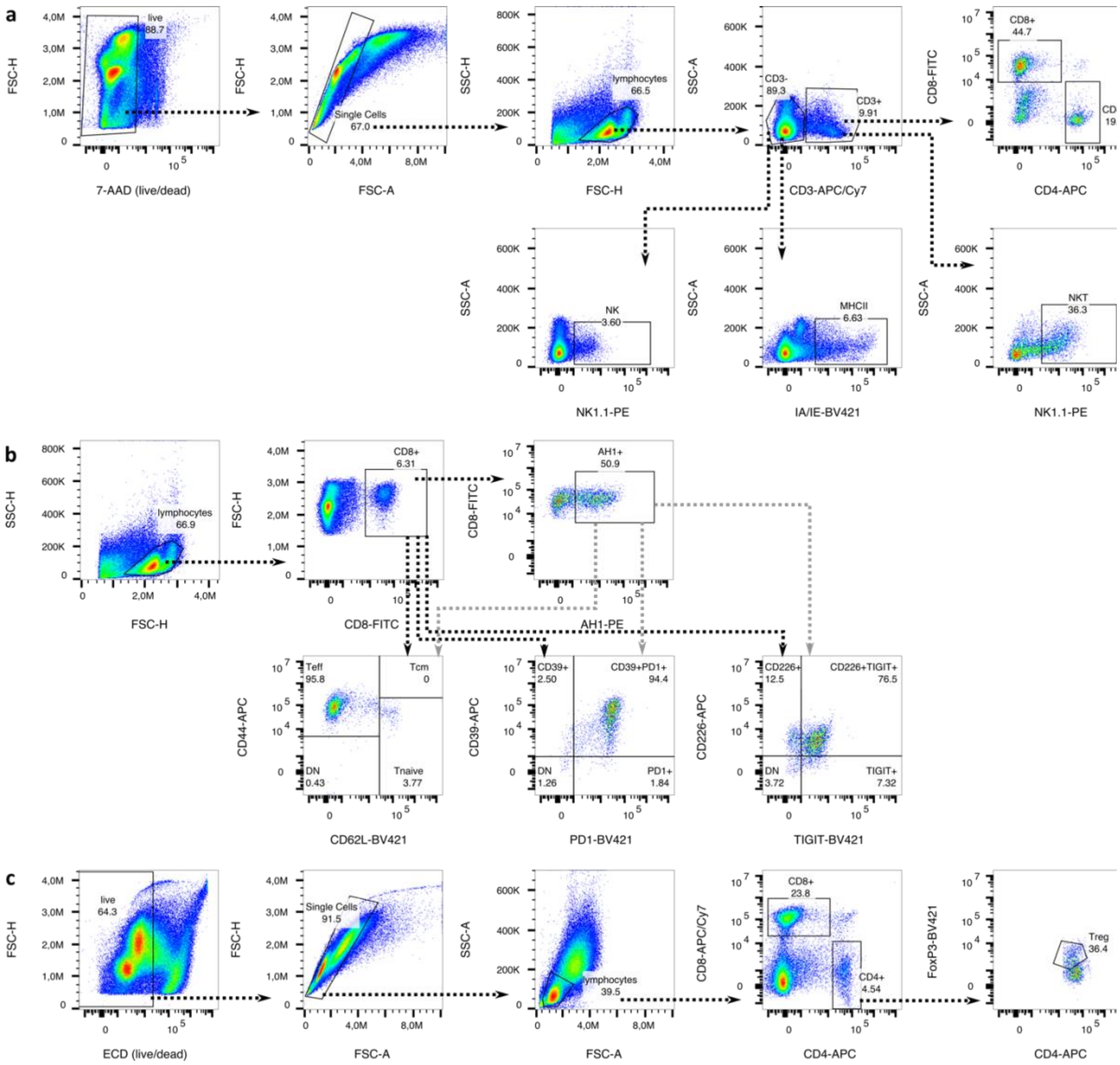


Supplementary Figure 6: Gating strategy for the analysis of tumor-infiltrating lymphocytes and TDLN (a) samples were first gated for live cells followed by a singlet gating and gating for lymphocytes by scattering. To quantify the cell subsets, cells were divided by the expression of CD3 and then further analyzed for expression of CD4, CD8, NK1.1 and MHC-II (IA/IE) (b) To analyze the phenotype of CD8+ T cells, the cells were gated for lymphocytes as shown in part a and then the same gates were applied to bulk CD8+ T cells and AH1-specific T cells. (d) Fixed samples were also first gated for live cells, singlets and lymphocytes before they were analyzed for the expression of CD8, CD4 and FoxP3. The gating strategy is shown for 1 representative tumor sample, but the same gates were applied to all samples.
